# Which are the central aspects of infant sleep? The dynamic of sleep composites across infancy

**DOI:** 10.1101/2020.10.26.354803

**Authors:** Sarah F. Schoch, Reto Huber, Malcolm Kohler, Salome Kurth

**Affiliations:** Department of Pulmonology, University Hospital Zurich, Zurich, CH; University of Zurich, Zurich, CH; Child Development Center, University Children’s Hospital Zurich, Zurich, CH; Department of Child and Adolescent Psychiatry and Psychotherapy, Psychiatric Hospital, University of Zurich, CH; Department of Psychology, University of Fribourg, Fribourg, CH

**Keywords:** actimetry, sleep assessment, maturation, sleep variables, variable selection

## Abstract

Sleep is ubiquitous during infancy and important for the well-being of both infant and parent. Therefore, there is large interest to characterize infant sleep with reliable tools, for example by means of combining actigraphy with 24-h-diaries. However, it is critical to select the right variables to characterize sleep. With a principal component analysis, we identified 5 underlying sleep composites from 48 commonly used sleep variables: *Sleep Night, Sleep Day, Sleep Activity, Sleep Timing and Sleep Variability.* These composites accurately reflect the known changes of sleep throughout infancy as *Sleep Day* (representing naps)*, Sleep Activity* (representing sleep efficiency and consolidation) and *Sleep Variability* (representing day-to-day stability) decrease across infancy, while *Sleep Night* (representing nighttime sleep) slightly increases and *Sleep Timing* becomes earlier with increasing age. Additionally, we uncover interesting dynamics between the sleep composites and demonstrate that infant sleep is not only highly variable between infants but also considerably dynamic within infants across time. Interestingly, *Sleep Day* is associated with behavioral development and therefore a potential marker for maturation. We recommend the use of sleep composites or of those specific single variables, which are solid representatives of the sleep composites for more reliable research.

## 1. Introduction

Why is sleeping the most common behavior of an infant in its first year of life [1]? Sleep fulfills an important function in development as the neurophysiology of sleep is linked to brain maturation, neural reorganization [2–4] as well as processes of learning and memory [5,6] [for an overview see 7]. However, besides the vital importance of sleep for the child, it also affects the quality of infant-parent bonds, as early periods with infant sleep problems have been linked to parental depression and stress [8,9]. Supporting healthy infant sleep can thus improve the wellbeing of the whole family.

Sleep-wake patterns are extensively diversified across infants – and vary to a much greater extent compared to any other period in life [1]. This inter-individual variability confounds the establishment of normative age-specific sleep values. Additionally, sleep is not a one-dimensional construct, but characterized by numerous dimensions of quantity, quality, timing or consolidation. While sleep undergoes drastic changes across infancy, not all sleep dimensions evolve at the same time or to the same degree. Possibly most recognizable is the alteration from sleep being distributed throughout the 24-hour-day (polyphasic sleep) to one primary sleep phase at nighttime (monophasic sleep) – a transition happening gradually from birth until about 5 years of age [1]. This transition involves a multitude of changes, as it affects not only the timing of sleep, but also its depth [10], and fragmentation [11]. Additionally, sleep quantity, measured as total sleep duration across 24 h, also decreases across the first year of life by ~8 minutes per month [12]. Notably, alongside the changes in sleep behavior, neurophysiology of sleep is reorganized and the composition of sleep states change across the first years of life: rapid eye movement (REM) sleep becomes less predominant and the electrophysiological characteristics typical for adult sleep (sleep spindles, slow waves) emerge [13,14].

Because of the ubiquity of sleep and its importance in early development, it is unsurprising that there is a large scientific interest in infant’s sleep. Researchers use both subjective methods (questionnaires on status-quo sleep, most widely used are the Brief Infant Sleep Questionnaire [BISQ] and 24-h-sleep-wake-diaries [15]) and objective methods (actigraphy [16], videosomnography [17] and polysomnography [18]). Each method has advantages and disadvantages [19]. There is only moderate agreement among the diverse methods, with larger discrepancies between questionnaires vs. objective data than between 24-h-diaries vs. objective data [20,21]. Subjective methods are costeffective and easy to administer to large populations. Yet, they are limited to items parents are aware of (e.g. sleep behavior but not sleep stages) and might be biased by parent’s perception. Furthermore, the selection of assessment method largely depends on the research question and available resources. Objective methods reduce subjective bias and comprehensively represent the different dimensions of sleep. Over the past 25 years, the combination of actigraphy with 24-h-diaries has emerged as the preferred method for many infant sleep investigations [22]. The advantage is its combination of objective and subjective methods which allows for the quantification of sleep in large populations and in natural environments, while being cost-effective [22,23]. However, issues remain regarding the standardization of actigraphy, especially in infants and young children [22,24]. One matter of improvement is the standardized reporting of methodological specifics, which is fundamental to this approach [23]. However, a remaining issue lies in the operationalization of sleep to capture its multiple dimensions accurately.

One current issue in investigating infant sleep is the selection of sleep variables. On one hand, there are several possible sleep domains and thus numerous sleep variables that can be calculated. On the other hand, the computation of these sleep variables is not standardized. The current situation leaves researchers to decide which sleep variable and which computations to choose [22]. For example, sleep duration is one of the most investigated sleep behaviors in infancy (reported in 82% of studies [12]). However, reports are based on different concepts, such as sleep duration computed across night-time only, sleep duration including 24 hours, duration of sleep with a split of day/night at a chosen clock time, or with clock times for day/night split that are individually assigned for each infant. This divergence is problematic, because it prevents comparability across studies [25]. It is also a likely source for lacking reproducibility. Additionally, researchers might rely on default variables from an automated analysis program, which is dubious if the research question demands more specificity. Using a large number of sleep variables to address the dimensions of sleep will likely increase false positives due to multiple testing [26]. Therefore, one should aim for a reduction of methodological complexity.

A novel and promising approach to handle the complexity of sleep dimensions was recently presented. Based on data of young children, Staples et al. proposed “sleep composites” that were combined from multiple commonly-used sleep variables. This approach reduces the dependence on single (often overlapping) sleep variables and increases the measurement stability [27]. A total of 4 sleep composites were discovered in both children and their mothers. These sleep composites contain the key dimensions of sleep; *Sleep Duration,* reflecting the quantity of sleep during the night; *Sleep Timing,* reflecting bedtimes and sleep onset times; *Sleep Variability,* reflecting day-to-day differences in sleep timing and duration; and *Sleep Activity,* reflecting movements and awakenings during the night. They also found that daytime sleep and sleep latency were separate constructs to these four sleep composites *(i.e.* loaded on their own composite). The identified sleep composites revealed higher consistency across different assessment timepoints compared to single sleep variables. A higher consistency is important to anchor sleep behaviors as reference in certain age periods, which is crucial, for example, to unravel the influence of early sleep variables on later regulatory, cognitive or emotional outcomes.

The goal of this study was to extend the approach of Staples et al., to an infant dataset to facilitate variable selection for future sleep studies. We included 48 single sleep variables, which thoroughly characterize the diverse dimensions of sleep, and performed a component analysis to identify the core infant sleep composites. We then examined the evolution of the sleep composites across repeated assessments throughout the first year of life, and tested for sex differences in the sleep composites. Additionally, we explored the stability of composites as well as the stability of the single sleep variables. To evaluate the relevance of sleep for development and to identify maturational markers we linked sleep composites to infant behavioral developmental scores.

## 2. Materials and Methods

### 2.1 Participants

152 healthy infants (69 female) in Switzerland participated in a longitudinal study on infant sleep and behavioral development. Of these, a subsample of 50 infants has been included in a previous investigation [24]. Caregivers and participants were recruited through maternity wards, midwifes, pediatricians, daycares, letters, social media, personal contacts and flyers distributed at universities, libraries, supermarkets, schools, family organizations and community centers. Participants were screened for study eligibility by means of an online questionnaire or telephone interview. Inclusion criteria for infants were good general health, being primarily breastfed at time of inclusion (*i.e.*, inclusion criterium of at least 50% of daily nutrition intake through breastfeeding at the first assessment at age 3 mo), vaginal birth (no cesarean section), and birth within 37-43 weeks of gestation. Parents were required to have good knowledge of German language.

Exclusion criteria for infants were disorders of the central nervous system, acute pediatric disorders, brain damage, chronic diseases as well as family background of narcolepsy, psychosis or bipolar disorder. Infants with birth weight below 2500 g, intake of medication affecting the sleepwake cycle, or antibiotics prior to the first assessment were also excluded.

Ethical approval was obtained from the *cantonal ethics committee* (BASEC 2016-00730) and study procedures were consistent with the declaration of Helsinki. Written parental consent was obtained after explanation of the study protocol and before enrollment.

### 2.2 Experimental design

We assessed 152 infants longitudinally at ages 3 mo, 6 mo and 12 mo. We scheduled assessments within a 1-month window around the target age, therefore actual age at start of assessment was between 2.43 – 3.39 mo, 5.42 – 6.28 mo and 11.47 – 12.26 mo.

We comprehensively quantified sleep-wake behavior for 11 continuous days. Ankle actigraphy and a 24-h-diary were simultaneously acquired during each of the three assessments, in alignment with our published recommendations for studying this age group [23]. GENEActiv movement sensors “actigraphs” (Activinsights Ltd, Kimbolton, UK, 43×40×13 mm, MEMS sensor, 16 g, 30 Hz Frequency recording resolution), which are sensitive for +/− 8 g range at 3.9 mg resolution, were attached to the infant’s left ankle in a modified sock (pocket sewn onto its side) or with a Tyvek paper strap. Parents were instructed to only remove the actigraph for bathing/swimming activities and to document any removal of the actigraph in the 24-h-diary. In the 24-h-diary (adapted from [21]) parents reported in 15-minute intervals about infant sleep and external movement occurring during infant sleep, e.g., sleeping in the parents arms, stroller, or baby sling etc. Further recorded parameters included feeding, crying episodes (> 15 minutes) and bed times (putting infant to bed in the evening and picking it up from the bed in the morning).

Additionally, in online questionnaires parents reported information on family background, health and demographics. Families received small gifts for their participation.

### 2.3 Behavioral development

Behavioral developmental status was assessed with the age-appropriate Ages and Stages questionnaire [28]. A *Collective Score*, represented by the sum of scores across five sub-domains (*Communication*, *Gross Motor*, *Fine Motor*, *Problem Solving* and *Personal Social*), was computed to quantify overall development. Additionally, we analyzed *Personal Social* and *Gross Motor* individually, because these subscales correlated with the well-validated testing battery Bayley Scales of Infant Development [29] and specifically also because these two sub-domains can indicate developmental delay [28,30]. Participants whose questionnaire was completed later than 1 week after the last day of the corresponding assessment were excluded from analysis, and missing data was imputed (section 2.4.2).

### 2.4 Sleep analysis

#### 2.4.1 Sleep–wake-behavior

Actigraph data was processed according to our standard protocols [24]. Binary data were extracted using GENEactiv PC Software (Version 3.1), imported into Matlab (R2016b) and converted to activity counts [31]. The latter included a 3-11 Hz bandpass filter and signal compression to 15 s bins. Acceleration data from the three movement axes was combined using sum of squares. The signal was then compiled to one data point per minute (analysis resolution). A published algorithm [32] was used to identify infant sleep and wake periods, and a 6-step modification [24] was applied to refine prediction for a better fit with the 24-h-diary. The first step of the modification (distinction between periods of high and low activity) was adjusted to use a threshold of ‘mean activity * 0.72’. Time periods without actigraphy information (*i.e.*, when the actigraph was not worn) were identified through the 24-h-diary or visual inspection (abrupt periods of no activity) and completed with information provided in the 24-h-diary.

#### 2.4.2 Handling of missing data

For some infants no sleep data was available for all timepoints: n = 2 at 3 mo (study enrollment at later age), n = 4 at 6 mo (n = 3 device failure, n = 1 parent withdrew from sleep assessment part of study) and n = 9 at 12 mo (n = 2 device failure, n = 3 participant attrition, n = 2 parent withdrew from sleep assessment part of study, n = 1 family moved away, n = 1 chronic sickness). Participants were instructed to collect actigraphy data for the duration of 11 continuous days (i.e., putting actigraph on before bedtime on the first day and removing it after getting up on the last day). Yet sickness and vacation of participants as well as device failure prevented the full 11-day recording in some cases (n = 10 at 3 mo, n = 28 at 6 mo, n = 28 at 12 mo). Further, in some instances the recording period was extended beyond the 11 days (e.g. because the original device was temporarily lost or parents recorded longer, n = 27 at 3 mo, n = 23 at 6 mo, n = 15 at 12 mo). Therefore, recordings with available data for both actigraphy and 24-h-diary lasted on average 10.76 ± 1.72 days: 11.13 ± 1.17 days at 3 mo, 10.60 ± 1.91 at 6 mo and 10.55 ± 1.93 at 12 mo. Additionally, single days were excluded if infants were either sick (except for common cold symptoms), or if the actigraph was removed for a longer time duration or if the fit between actigraphy-based data and 24-h-diary was poor (see Table A1).

#### 2.4.3 Calculation of sleep variables

To capture the multitude of dimensions of infants’ sleep, we calculated 48 sleep variables of interest, based on previous definitions [22,27,33] (Table 1). 3 valid recording days of actimetry were set as minimum to compute sleep variables in each participant. For variability variables a minimum of 5 valid recording days was required. All calculated sleep variables (except variability variables which were standard deviations across days) were averaged across all valid recorded days. After calculating sleep variables, additional exclusions were performed: for time zone change of > 1h less than 1 week before the recording (n = 1 at 12 mo), for medication affecting sleep (n = 2 at 3 mo) and for medical problems (n = 1 at 6 mo, n = 2 at 12 mo) or psychological trauma was experienced (n = 1 at 12 mo).

**Table 1.** Definition and descriptive statistics at 3,6 and 12 months of the 48 infant sleep variables based on 24-h-diary and actigraphy that entered the principal component analysis. Bedtime and Get up Time and their variability variables are based on parent report in 24-h-diary, all other variables are based on actigraphy (with adjustments from diaries as reported in 2.4.1). Mean ± SD (Minimum – Maximum)

| Variable Name | 3 Months | 6 Months | 12 Months |
| --- | --- | --- | --- |
| (1) *Bedtime* (clock time in min)<br>Parent-reported time in the 24-h-diary of putting the child to bed. For missing values, the first minute of reported sleep was used. If bedtime exceeded midnight 1440 was added. | 1273.71 ± 75.38<br>(1116 - 1479) | 1226.59 ± 69.18<br>(1105 - 1420.8) | 1220.78 ± 53.46<br>(1120 - 1416) |
| (2) *Variability of Bedtime* (SD)<br>Standard deviation of *Bedtime* across recording days | 43.5 ± 21.64<br>(0 - 118.39) | 32.6 ± 17.76<br>(0 - 87.13) | 28.88 ± 16.83<br>(3.35 - 91.41) |
| (3) *Get up Time* (clock time in min)<br>Parent reported time in the 24-h-diary of getting out of bed in the morning. For missing values, the last minute of reported sleep was used. | 472.88 ± 52.06<br>(361.5 - 612) | 443.84 ± 48.89<br>(335.5 - 577.7) | 437.93 ± 43.41<br>(323.33 - 625) |
| (4) *Variability of Get up Time* (SD)<br>Standard deviation of *Get up Time* across recording days | 42.02 ± 16.44<br>(6.35 - 96.17) | 36.35 ± 16.26<br>(0 - 109.2) | 35.8 ± 16.93<br>(6.12 - 99.82) |
| (5) *Sleep Onset* (clock time in min)<br>Following *Bedtime*, the first minute asleep of at least 10 minutes of consecutive sleep. If asleep at *Bedtime*, the first minute asleep before *Bedtime* was chosen. | 1257.94 ± 68.08<br>(1127.17 - 1473.3) | 1228.59 ± 65.87<br>(1112.33 - 1449.4) | 1228.74 ± 54.7<br>(1125.5 - 1423.6) |
| (6) *Variability of Sleep Onset* (SD)<br>Standard deviation of *Sleep Onset* across recording days | 51.87 ± 23.83<br>(7.75 - 157.33) | 37.79 ± 19.07<br>(6.33 - 109.24) | 34.28 ± 17.2<br>(4.7 - 79.86) |
| (7) *Sleep Latency* (min)<br>Duration in minutes between *Bedtime* and *Sleep Onset*, set to 0 if *Sleep Onset* is before *Bedtime* | 7.79 ± 7.97<br>(0 - 42) | 11.29 ± 8.3<br>(0 - 45.5) | 10.94 ± 8.2<br>(0 - 38.38) |
| (8) *Variability of Sleep Latency* (SD)<br>Standard deviation of *Sleep Latency* across recording days | 10.29 ± 9.17<br>(0 - 56.64) | 11.43 ± 7.61<br>(0 - 44.91) | 10.26 ± 7.14<br>(0 - 40.27) |
| (9) *Sleep Offset* (clock time in min)<br>Last minute asleep of at least 10 consecutive minutes asleep before *Get up Time* or if asleep at *Get up Time* last minute asleep after *Get up Time* | 470.93 ± 53.13<br>(354.8 - 622.86) | 437.71 ± 49.41<br>(324.5 - 583.38) | 437.96 ± 46.6<br>(316.67 - 615.5) |
| (10) *Variability of Sleep Offset* (SD)<br>Standard deviation of *Sleep Offset* across recording days | 48.06 ± 18.51<br>(15.82 - 114.17) | 39.74 ± 16.89<br>(16.51 - 123.17) | 38.66 ± 20.5<br>(12.02 - 164.83) |
| (11) *Midsleep* (clock time in min)<br>Midpoint between *Sleep Onset* and *Sleep Offset* | 143.93 ± 53.02<br>(32.65 - 298.7) | 112.88 ± 51.71<br>(13.7 - 293.81) | 114.03 ± 46.09<br>(13.5 - 278.2) |
| (12) *Variability of Midsleep* (SD)<br>Standard Deviation of *Midsleep* across recording days | 37.62 ± 13.62<br>(11.95 - 78.54) | 28.37 ± 11.24<br>(8.86 - 64.22) | 27.33 ± 13.46<br>(7.7 - 101.79) |
| (13) *Sleep Opportunity* (min)<br>Time between *Bedtime* and *Get Up Time* (unless asleep at either of these times, in which case *Sleep Onset/Sleep Offset* was used) | 662.46 ± 62.8<br>(494.2 - 840) | 670.89 ± 54.38<br>(558.11 - 820.67) | 666.4 ± 42.51<br>(578 - 738) |
| (14) *Variability of Sleep Opportunity* (SD)<br>Standard deviation of *Sleep Opportunity* across recording days | 62.85 ± 23.14<br>(16.83 - 124.55) | 47.29 ± 20.18<br>(12.31 - 129.93) | 45.32 ± 23.34<br>(14.51 - 163.62) |
| (15) *Sleep Period (min)*<br>Time between *Sleep Onset* and *Sleep Offset* | 651.87 ± 58.05<br>(488.9 - 796) | 651.52 ± 50.44<br>(543.56 - 821.78) | 650.94 ± 44.7<br>(547 - 735.75) |
| (16) *Variability of Sleep Period (SD)* | 67.09 ± 22.21<br>(21.19 - 127.12) | 51.83 ± 21.73<br>(16.36 - 144.75) | 49.85 ± 23.43<br>(11.27 - 163.32) |
| (17) *Total Sleep Time (min)*<br>Minutes scored 'Sleep' within *Sleep Period* | 573.63 ± 58.25<br>(421.33 - 709.44) | 605.23 ± 47.38<br>(492.44 - 728.75) | 627.3 ± 51.29<br>(488.5 - 717.1) |
| (18) *Variability of Total Sleep Time (SD)*<br>Standard deviation of *Total Sleep Time* across recording days | 53.47 ± 18.91<br>(18.62 - 108.3) | 45.98 ± 16.04<br>(10.15 - 91.2) | 46.33 ± 19.44<br>(11.61 - 140.08) |
| (19) *Sleep Efficiency (%)*<br>(*Total Sleep Time*)/(*Sleep Opportunity*) * 100 | 87.83 ± 5.4<br>(69.37 - 99.5) | 90.67 ± 4.08<br>(80.55 - 99.21) | 94.25 ± 3.52<br>(84.06 - 99.64) |
| (20) *Variability of Sleep Efficiency (SD)*<br>Standard deviation of *Sleep Efficiency* across recording days | 5.46 ± 2.35<br>(1.84 - 18.22) | 4.59 ± 1.95<br>(1.57 - 11.97) | 3.57 ± 1.9<br>(0.45 - 12.94) |
| (21) *Wake after Sleep Onset (min)*<br>Minutes scored 'Wake' in *Sleep Period* | 69.04 ± 32.15<br>(13.7 - 197.75) | 44.73 ± 24.55<br>(1.86 - 121.17) | 22.31 ± 17.02<br>(0 - 78.22) |
| (22) *Variability of Wake after Sleep Onset (SD)*<br>Standard deviation of *Wake after Sleep Onset* across recording days | 32.72 ± 12.52<br>(11.3 - 79.02) | 26.63 ± 13.46<br>(4.26 - 86.03) | 18.8 ± 10.68<br>(1.9 - 57.17) |
| (23) *Longest Nocturnal Wake (min)*<br>Longest period scored 'Wake' followed by at least 15 mins scored 'Sleep' in *Sleep Period* | 31.87 ± 14.5<br>(7.6 - 93.33) | 24.37 ± 13.19<br>(1.86 - 75.89) | 13.74 ± 9.65<br>(0 - 56.67) |
| (24) *Variability of Longest Nocturnal Wake (SD)*<br>Standard deviation of *Longest Nocturnal Wake* across recording days | 16.43 ± 7.32<br>(3.1 - 39.7) | 17.09 ± 10.2<br>(3.14 - 48.35) | 12.04 ± 8.78<br>(0 - 60.19) |
| (25) *Nocturnal Wake Frequency per Hour (wakings/hour)*<br>(Number of Nocturnal Wake Periods in *Sleep Period*)/*Sleep Period* | 0.34 ± 0.11<br>(0.1 - 0.61) | 0.23 ± 0.1<br>(0.02 - 0.49) | 0.14 ± 0.09<br>(0 - 0.46) |
| (26) *Variability of Nocturnal Wake Frequency per Hour (SD)*<br>Standard deviation of *Nocturnal Wake Frequency per Hour* across recording days | 0.12 ± 0.04<br>(0.03 - 0.27) | 0.11 ± 0.04<br>(0.03 - 0.22) | 0.1 ± 0.04<br>(0 - 0.24) |
| (27) *Variability of Activity level (SD)*<br>Standard deviation of activity per minute in *Sleep Period* | 168.64 ± 61.51<br>(40.69 - 356.73) | 178.91 ± 79.21<br>(49.92 - 478.14) | 108.5 ± 50.86<br>(33.64 - 336.46) |
| (28) *Percent Active Epochs (ratio)*<br>(Minutes of epochs with non-zero activity in *Sleep Period*)/*Sleep Period* | 0.3 ± 0.05<br>(0.14 - 0.4) | 0.24 ± 0.04<br>(0.13 - 0.34) | 0.23 ± 0.03<br>(0.14 - 0.32) |
| (29) *Variability Percent Active Epochs (SD)*<br>Standard deviation of *Percent Active Epochs* across recording days | 0.04 ± 0.01<br>(0.02 - 0.09) | 0.04 ± 0.02<br>(0.01 - 0.1) | 0.04 ± 0.01<br>(0.01 - 0.12) |
| (30) *Longest Sleep (min)*<br>Longest continuous period scored as 'Sleep' | 292.19 ± 90.16<br>(139 - 580.29) | 339.21 ± 99.18<br>(158.75 - 632.17) | 458.24 ± 126.82<br>(166.89 - 706.44) |
| (31) *Variability of Longest Sleep (SD)*<br>Standard deviation of *Longest Sleep* across recording days | 82.75 ± 36.25<br>(21.04 - 190.74) | 104.82 ± 39.39<br>(23.02 - 222.59) | 127.41 ± 50.7<br>(14.21 - 245.62) |
| (32) *Longest Wake (min)*<br>Longest continuous period scored as 'Wake' | 162.44 ± 27.74<br>(101.11 - 292.29) | 212.13 ± 32.94<br>(139.33 - 348) | 293.14 ± 40.25<br>(195.33 - 402) |
| (33) *Variability of Longest Wake (SD)*<br>Standard deviation of *Longest Wake* across recording days | 40.07 ± 18<br>(11.65 - 111.57) | 50.11 ± 22.36<br>(9.07 - 109.98) | 65.91 ± 22.99<br>(18.63 - 150.2) |
| (34) *Nap Counter*<br>Number of daytime sleep periods exceeding 20 minutes between *Sleep Offset* and *Sleep Onset* | 4.06 ± 0.77<br>(2 - 6.25) | 3.2 ± 0.59<br>(1.38 - 4.56) | 2.07 ± 0.55<br>(0.67 - 3.57) |
| (35) *Variability Nap counter (SD)*<br>Standard deviation of *Nap counter* across recording days | 1.1 ± 0.31<br>(0.38 - 2.32) | 0.84 ± 0.3<br>(0 - 1.72) | 0.74 ± 0.27<br>(0 - 1.9) |
| (36) *Sleep after Wake Onset (min)*<br>Minutes scored Sleep between *Sleep Offset* and *Sleep Onset* | 247.56 ± 53.3<br>(123 - 382.88) | 179.03 ± 37.26<br>(95.6 - 298.43) | 142.54 ± 38.17<br>(70.5 - 282.63) |
| (37) *Variability Sleep after Wake Onset (SD)*<br>Standard deviation of *Sleep after Wake Onset* across recording days | 61.92 ± 20.6<br>(19.22 - 154.89) | 45.27 ± 16.89<br>(15.49 - 112.18) | 43.76 ± 17.8<br>(10.94 - 140.06) |
| (38) *Sleep Duration 24 h (min)*<br>Minutes scored 'Sleep' across 24 h | 822.19 ± 55.68<br>(672.86 - 975.44) | 783.15 ± 44.45<br>(654 - 922.33) | 767.64 ± 45.34<br>(609.67 - 867.43) |
| (39) *Variability of Sleep Duration 24 h (SD)*<br>Standard deviation of *Sleep Duration 24 h* across recording days | 66.34 ± 19.58<br>(28.03 - 123.61) | 58.77 ± 19.91<br>(24.73 - 121.56) | 54.17 ± 18.89<br>(20.77 - 127.09) |
| (40) *Sleep Duration Day (min)*<br>Minutes scored 'Sleep' between 7 am and 7 pm | 278.59 ± 44.98<br>(158.7 - 396) | 203.46 ± 39.66<br>(115.88 - 345.13) | 165.83 ± 41.31<br>(81.33 - 298.56) |
| (41) *Variability Sleep Duration Day (SD)*<br>Standard deviation of *Sleep Duration Day* across recording days | 53.42 ± 16.05<br>(17.33 - 125.21) | 43.44 ± 14.42<br>(15.79 - 83.25) | 43.2 ± 12.34<br>(21.21 - 76.36) |
| (42) *Sleep Duration Night (min)*<br>Minutes scored 'Sleep' between 7 pm and 7 am | 548.43 ± 45.54<br>(407.22 - 644.44) | 579.11 ± 49.17<br>(426.75 - 672.11) | 602.1 ± 47.78<br>(467.89 - 699.11) |
| (43) *Variability of Sleep Duration Night (SD)*<br>Standard deviation of *Sleep Duration Night* across recording days | 44.78 ± 16.37<br>(12.78 - 92.5) | 40.95 ± 14.53<br>(12.34 - 103.61) | 35.51 ± 13.72<br>(7.31 - 72.05) |
| (44) % *Sleep Duration Night (ratio)*<br>(*Sleep Duration Night*) / (*Sleep Duration 24h*) | 0.67 ± 0.05<br>(0.55 - 0.84) | 0.74 ± 0.05<br>(0.57 - 0.85) | 0.78 ± 0.05<br>(0.61 - 0.88) |
| (45) *Variability % Sleep Duration Night (SD)*<br>Standard deviation of % *Sleep Duration Night* across recording days | 0.05 ± 0.01<br>(0.02 - 0.09) | 0.05 ± 0.01<br>(0.02 - 0.11) | 0.05 ± 0.01<br>(0.02 - 0.1) |
| (46) *Sleep Regularity Index Whole Day (ratio)*<br>The probability of being in the same state (Sleep or Wake) computed for each minute, averaged across one day, and then across all recording days. Represented with ratio (0-1) ('Sleep'/'Wake), where 1 reflects the exact same rhythm every day | 0.77 ± 0.03<br>(0.66 - 0.84) | 0.82 ± 0.03<br>(0.74 - 0.89) | 0.87 ± 0.03<br>(0.76 - 0.95) |
| (47) *Sleep Regularity Index Day (ratio)*<br>*Sleep Regularity Index* for the clock times from 7 am and 7 pm | 0.7 ± 0.04<br>(0.62 - 0.86) | 0.76 ± 0.04<br>(0.64 - 0.91) | 0.81 ± 0.04<br>(0.7 - 0.92) |
| (48) *Sleep Regularity Index Night (ratio)*<br>*Sleep Regularity Index* for the clock times from 7 pm and 7 am | 0.84 ± 0.05<br>(0.66 - 0.96) | 0.88 ± 0.05<br>(0.62 - 0.97) | 0.92 ± 0.04<br>(0.79 - 0.99) |

#### 2.4.4 Data Imputation

Subsequent analyses were done in R (version 3.5.0) [34] and R studio (version 1.1.463) [35], with several packages for data handling (*tidyr, eeptools, reshape, dplyr, lubridate, phyloseq, VIM, margrittr, chron, kableExtra, knitr, qwraps2*) and plotting (*corrplot, ggplot2, lattice, ggfortify, sjPlot, cowplot*) [36–53]. Missing and excluded data was imputed using multiple imputation in the *mice* package [54] and additional functions from *miceadds, MKmisc* and *micemd* package [55–57]. Missing data ranged from 0% to 22.32% per variable. The dataset used for imputation included all sleep variables and several demographic variables (see Appendix A). All numerical variables were predicted using the method “2l.pmm”, using the participant ID as grouping variable and assessment age (3/6/12 months) as slope. Binary variables were predicted using the method “logreg” and categorical variables were predicted using either “polyreg” or “polyr”. Two-level structure was not included in binary and categorical variable prediction. 100 imputations were run with 100 iterations each using 5 cores (20 imputations per core). Data quality of the imputations were visually controlled with density plots (observed vs. imputed values) and line plots for convergence of the iterations. The reported method and prediction matrix were chosen due to best fit of the density plot.

#### 2.4.5 Sleep Composites

We used an integrative and data-driven approach to congregate the 48 infant sleep variables (such as *Total Sleep Time*, see Table 1 for full description) to the core composite scores, inspired by an approach in young children [58]. We applied a principal component analysis (PCA) with *promax rotation* (*psych* package [59]) across all participants and all assessment timepoints. Because we included more variables than Staples et al., we examined the best solution with scree and parallel plots as well as the interpretability of the resulting composites. Therefore, we ended up choosing a 5-component solution. We removed single sleep variables with absolute factor loadings below 0.512 as recommended for sample sizes exceeding 100 [60]. This led to the exclusion of 14 variables (see Table 2). Additionally, we excluded *Sleep Duration 24 h* (min, minutes scored ‘Sleep’ across 24 hours) for interpretability (details below). In total, 33 variables were included in the final PCA solution, with 3 to 10 single sleep variables assigned to each sleep composite (Table 2).

**Table 2.** Single sleep variables and PCA solution with oblique rotation (promax rotation). The numbers in parentheses link the single sleep variables to their explanation in Table 1. Values in bold indicate the strongest loading. 14 variables were excluded due to low loading (< 0.512) on any of the sleep composites. These were (7) *Sleep Latency* (min), (8) *Variability of Sleep Latency* (SD), (20) *Variability of Sleep Efficiency,* (26) *Variability of Nocturnal Wake Frequency per Hour* (SD), (29) *Variability Percent Active Epochs* (SD), (31) *Variability of Longest Sleep* (SD), (35) *Variability Nap counter* (SD), (37) *Variability Sleep after Wake Onset* (SD), (39) *Variability of Sleep Duration 24 h* (SD), (41) *Variability Sleep Duration Day* (SD), (42) *Sleep Duration Night* (min), (43) *Variability of Sleep Duration Night* (SD), (45) *Variability % Sleep Duration Night* (SD), (46) *Sleep Regularity Index Whole Day* (Ratio).

| Variables | Sleep Activity | Sleep Variability | Sleep Day | Sleep Timing | Sleep Night |
| --- | --- | --- | --- | --- | --- |
| (19) *Sleep Efficiency* (%) | **-0.89** | 0.05 | 0.00 | -0.01 | -0.02 |
| (23) *Longest Nocturnal Wake* (min) | **0.88** | 0.04 | 0.05 | 0.02 | 0.19 |
| (21) *Wake after Sleep Onset* (min) | **0.86** | -0.01 | -0.11 | 0.00 | 0.16 |
| (25) *Nocturnal Wake Frequency per Hour* (wakings/hour) | **0.79** | -0.13 | -0.14 | 0.03 | -0.13 |
| (30) *Longest Sleep* (min) | **-0.77** | 0.09 | 0.04 | -0.03 | 0.15 |
| (27) *Variability of Activity level* (SD) | **0.76** | -0.05 | 0.10 | 0.01 | 0.02 |
| (22) *Variability of Wake after Sleep Onset* (SD) | **0.72** | 0.09 | 0.10 | 0.01 | 0.05 |
| (24) *Variability of Longest Nocturnal Wake* (SD) | **0.70** | 0.07 | 0.28 | -0.05 | 0.02 |
| (48) *Sleep Regularity Index Night* (ratio) | **-0.69** | -0.18 | 0.17 | 0.04 | 0.09 |
| (28) *Percent Active Epochs* (ratio) | **0.58** | -0.08 | -0.28 | 0.01 | 0.12 |
| (14) *Variability of Sleep Opportunity* (SD) | -0.05 | **0.87** | -0.07 | -0.08 | 0.04 |
| (16) *Variability of Sleep Period* (SD) | 0.04 | **0.86** | 0.03 | -0.11 | -0.07 |
| (18) *Variability of Total Sleep Time* (SD) | -0.06 | **0.73** | 0.10 | 0.00 | 0.00 |
| (10) *Variability of Sleep Offset* (SD) | -0.09 | **0.73** | -0.05 | 0.02 | 0.11 |
| (12) *Variability of Midsleep* (SD) | 0.00 | **0.73** | -0.08 | -0.01 | -0.05 |
| (4) *Variability of Get up Time* (SD) | -0.13 | **0.72** | 0.02 | 0.14 | 0.19 |
| (6) *Variability of Sleep Onset* (SD) | 0.10 | **0.64** | -0.04 | 0.06 | -0.10 |
| (2) *Variability of Bedtime* (SD) | 0.20 | **0.58** | 0.05 | -0.02 | -0.11 |
| (32) *Longest Wake* (min) | -0.10 | 0.04 | **-0.92** | 0.11 | -0.17 |
| (34) *Nap Counter* | 0.03 | -0.06 | **0.86** | -0.12 | -0.14 |
| (36) *Sleep after Wake Onset* (min) | 0.00 | 0.01 | **0.82** | -0.11 | -0.26 |
| (47) *Sleep Regularity Index Day* (ratio) | -0.02 | -0.24 | **-0.76** | 0.07 | -0.02 |
| (40) *Sleep Duration Day* (min) | -0.01 | 0.11 | **0.72** | 0.32 | 0.05 |
| (33) *Variability of Longest Wake* (SD) | 0.06 | 0.12 | **-0.68** | 0.03 | -0.21 |
| (44) *% Sleep Duration Night* (ratio) | -0.12 | -0.05 | **-0.56** | -0.39 | 0.12 |
| (9) *Sleep Offset* (clock time in min) | 0.03 | -0.01 | 0.03 | **1.01** | 0.32 |
| (3) *Get up Time* (clock time in min) | 0.08 | -0.03 | 0.01 | **0.97** | 0.37 |
| (11) *Midsleep* (clock time in min) | -0.02 | -0.01 | 0.08 | **0.93** | -0.18 |
| (5) *Sleep Onset* (clock time in min) | -0.05 | -0.04 | 0.08 | **0.76** | -0.52 |
| (1) *Bedtime* (clock time in min) | -0.07 | 0.00 | -0.04 | **0.68** | -0.49 |
| (15) *Sleep Period* (min) | 0.11 | 0.03 | -0.01 | 0.12 | **0.99** |
| (13) *Sleep Opportunity* (min) | 0.18 | 0.00 | 0.05 | 0.08 | **0.96** |
| (17) *Total Sleep Time* (min) | -0.40 | -0.03 | 0.04 | 0.06 | **0.78** |
| Proportion of Variance explained | **0.19** | **0.14** | **0.14** | **0.13** | **0.11** |

Each subsequent model was run with all 100 imputations of the PCA-derived scores for each participant and sleep composite (unweighted average of the highest loadings). All results were pooled across all 100 models. To evaluate effects of age and sex we used linear regression models. With the *corrplot* package we examined correlations between sleep composites and assessment time points using Spearman correlation coefficients. Bonferroni correction was applied to address multiple comparison issues. To test the stability of effects across development, the range of each infant’s percentiles across all assessment timepoints was evaluated (within-subject stability). The stability of composites vs single variables was evaluated using paired t-tests.

Associations of sleep composites with behavioral outcomes were identified based on longitudinal multilevel models using the *lme4* package and by including participant ID for the intercepts and timepoint as slope. Covariates were exact age and sex, and predictors were the 5 sleep composites. Values were considered outliers if they exceeded 1.5 times the interquartile range below the 1st quartile or above the 3^rd^ quartile. Reported statistics include outliers, but any changes in significance due to exclusions of outliers are mentioned specifically. Significance level was set to below 0.05.

## 3. Results

### 3.1 Five principal components express all infant sleep variables: infant sleep composites

We achieved reduction of complexity of infant sleep variables by determining 5 core sleep composites. The relationship of each of the 48 original sleep variables with the sleep composites is represented as “loadings” (Table 2). The five sleep composites explain a total of 71% of the variances, which yields a diagonal fit of 0.98. This revealed:

- *Sleep Activity* – Larger values generally reflect more movements and more awakenings during the night as well as less regularity of awakenings. The most representative (i.e. with highest loading) single variable were *Sleep Efficiency* (negative) or *Longest Nocturnal Wake* (positive).
- *Sleep Timing* – Increased values generally reflect later clock time of bed times and sleep times. The most representative single variable was *Sleep Offset*.
- *Sleep Night* – Larger values indicate longer nighttime sleep opportunity and longer nighttime sleep duration. The most representative single variable was *Sleep Period*.
- *Sleep Day* – Larger values refer to longer daytime sleep duration, more daytime naps, and lower regularity in daytime sleep. The most representative variables were *Longest Wake* (negatively) or *Nap Counter* (positively).
- *Sleep Variability* – Larger values identify increased variability from day-to-day (standard deviation) within *Sleep Timing* and *Sleep Night*. The most representative single variable was *Variability of Sleep Opportunity*.

Interestingly *24 h Sleep Duration* showed the highest loading on the *Sleep Day* composite, meaning it was more related to *Sleep Day* than *Sleep Night*. This finding supports particularly the tight link between naps and total sleep duration. However, to make the interpretation of the *Sleep Day* composite easier, we removed this variable from subsequent analyses.

### 3.2 Sleep composites accurately reflect sleep maturation across infancy

To ensure that the sleep composites accurately reflect the maturation of sleep patterns in infancy, we examined changes in the sleep composites across age. As expected, *Sleep Activity, Sleep Day, Sleep Timing* and *Sleep Variability* all decreased with age (*Sleep Activity* t_(434.25)_ = −14.59, p < 0.001, *Sleep Day* (413.53) = −25.09, p < 0.001, *Sleep Timing* t_(426.65)_ = −5.78, p < 0.001, *Sleep Variability* t_(423.45)_ = −6.13, p < 0.001). In other words, in comparison to younger age, older infants showed lower activity at night and woke up less frequently (b = −0.15 per month older), slept less often and also shorter during the day (b = – 0.21), went to sleep earlier at night and woke up earlier in the morning (b = −0.07) and were more consistent in their sleep timing and nighttime sleep duration (b = −0.08). *Sleep Night* on the other hand, slightly increased with age (t_421.30)_ = 2.59, p = 0.01), indicating that older infants slept more at night (b = 0.03). Therefore, sleep composites capture the sleep maturation in infancy well. Moreover, within the same models we could observe sex differences, specifically in *Sleep Activity* (female vs male t_(431.04)_ = −3.84, p < 0.001) and *Sleep Variability (*t_(434.79)_ = −1.88, p = 0.06), yet the latter was significant only after exclusion of outliers (t_(413.77)_ = −2.21, p = 0.03). Girls showed lower nightly activity and reduced wakings (b = −0.29 for female) and were more consistent in their sleep routine (with outliers b = −0.17/ without outliers b = −0.19 for female). No sex differences were detected in the other sleep composites (p > 0.05).

### 3.3 Strong correlations between the sleep composites

Next, we investigated the interrelationships between the sleep composites. Notably, each sleep composite correlated significantly with all other sleep composites, indicating that while sleep is a multidimensional construct, the different dimensions are tightly intertwined (all p < 0.001; Figure 1). Interestingly, the strongest positive correlation was found between *Sleep Activity* and *Sleep Day* (r_s_ = 0.50, p < 0.001). Higher activity at night was associated with more sleep during the day. Surprisingly, this association was stronger than the association of *Sleep Activity* and *Sleep Night*. As expected, a strong positive correlation was found between *Sleep Timing* and *Sleep Variability* (r_s_ = 0.48, p < 0.001), i.e. the later the sleep timing, the higher the *Sleep Variability.* The strongest negative correlations were found between *Sleep Day* and *Sleep Night* (r_s_ = −0.26, p < 0.001) with infants that slept more during the day sleeping less at night. A strong negative association was also found for *Sleep Night* with *Sleep Timing* (r_s_ = −0.25, p < 0.001), such that infants with later sleep times had less nighttime sleep. In sum, even though the approach clearly identified 5 core sleep composite of infant sleep, those composites are also highly interrelated with each other.

**Figure 1.**
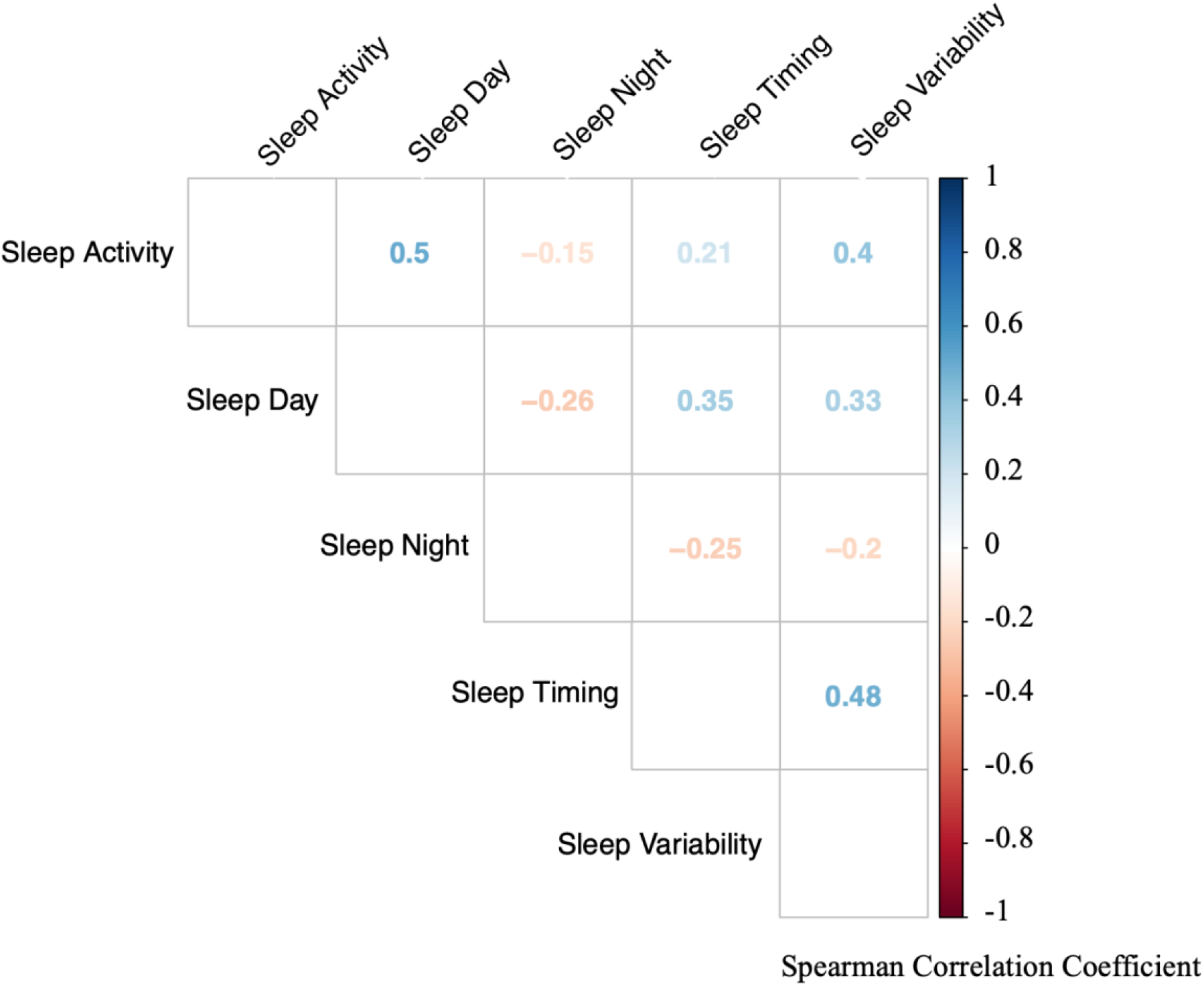
Correlations between the infant sleep composites based on all assessment timepoints. Each sleep composite is significantly associated with all other composites (all p < 0.001). Colors indicate strength of correlation (red = negative correlations, blue = positive correlations). Numbers indicate spearman correlation coefficient (rs).

### 3.3 Stability of sleep composites

To investigate the stability of sleep composites across the first infant year, we examined correlation coefficients between all assessment time points of each sleep composite (Table 3). Most sleep composites significantly correlated between the adjacent time points (3 *vs.* 6 or 6 *vs.* 12 months). Only *Sleep Timing* was also significantly correlated between 3 and 12 months. While *Sleep Variability* and *Sleep Night* were significantly correlated when outliers were removed, this correlation was low, suggesting no stability (R^2^ = 0.07). To understand the dynamics, we calculated the within-subject stability, *i.e.* consistency of the position of each subject in relation to all other participants. On average children had a maximum change of 29% for *Sleep Timing,* 38% for *Sleep Night,* 43% for *Sleep Variability* and *Sleep Day* and 45% for *Sleep Activity* from 3 – 12 months (values from one imputation). This suggests that although most sleep behaviors are stable in the short term, they are dynamic across the first year of infancy.

**Table 3.** Spearman Correlation Coefficients (rs) of Sleep Composites across assessment time points. Significant correlations are presented in bold (Bonferroni corrected p-value below 0.0033). The correlation marked with * are significant upon exclusion of outliers: *Sleep Variability* r_s_ = 0.26, p = 0.002, *Sleep Night* r_s_ = 0.26, p = 0.001). Composites are most stable across adjacent time points, but only *Sleep Timing* is stable across the entire first year.

| Sleep Composite | Correlation 3 vs. 6 Months |  | Correlation 6 vs. 12 Months |  | Correlation 3 vs. 12 Months |  |
| --- | --- | --- | --- | --- | --- | --- |
| | $r_s$ | p | $r_s$ | p | $r_s$ | p |
| Sleep Activity | **0.29** | **< 0.001** | 0.21 | 0.01 | 0.15 | 0.07 |
| Sleep Day | 0.25 | 0.004 | **0.29** | **< 0.001** | 0.11 | 0.21 |
| Sleep Night | **0.53** | **< 0.001** | **0.45** | **< 0.001** | 0.24* | 0.004 |
| Sleep Timing | **0.68** | **< 0.001** | **0.58** | **< 0.001** | **0.55** | **< 0.001** |
| Sleep Variability | **0.28** | **< 0.001** | **0.38** | **< 0.001** | 0.23* | 0.007 |

### 3.4 Stability of sleep composites vs single sleep variables

Subsequently we tested whether the sleep composites were more stable across the assessment timepoints compared to the stability of single sleep variables, as observed in young children and adults [27]. We used the within-subject stability and compared it between single and composite variables. We tested this within-subject stability in *Sleep Timing* (the most stable variable) and in *Sleep Activity,* (the least stable variable). Within-subject stability was also computed for the single sleep variables that loaded the highest and lowest on both sleep composites: *Sleep Offset* and *Bedtime* as well as *Longest Nocturnal Wake* and *Percent Active Epochs.* An exemplary comparison of one imputation and 6 random participants is shown in Figure 2. There was no significant difference between the within-subject stability of *Sleep Activity* and *Longest Nocturnal Wake* (t_(115.69)_ = −0.17, p = 0.86) nor between the within-subject stability of *Sleep Activity* and *Percent Active Epochs* (t_134.03)_ = 0.10, p = 0.92), indicating no advantage in within-subject stability in the sleep composite as compared to within-subject stability in single sleep variables. In other words, infants showed variable sleep behavior no matter how it was quantified. Similarly, there was no significant difference between the within-subject stability of *Sleep Timing* and *Bedtime* (t_124.88)_ = 0.40, p = 0.69). Contrastingly, *Sleep Timing* showed higher within-subject stability as compared to *Sleep Offset* (t_(106.64)_ = 3.20, p = 0.002). Thus, overall, we cannot confirm higher within-subject stability across the first year of life in sleep composites versus within-subject stability in single sleep variables.

**Figure 2.**
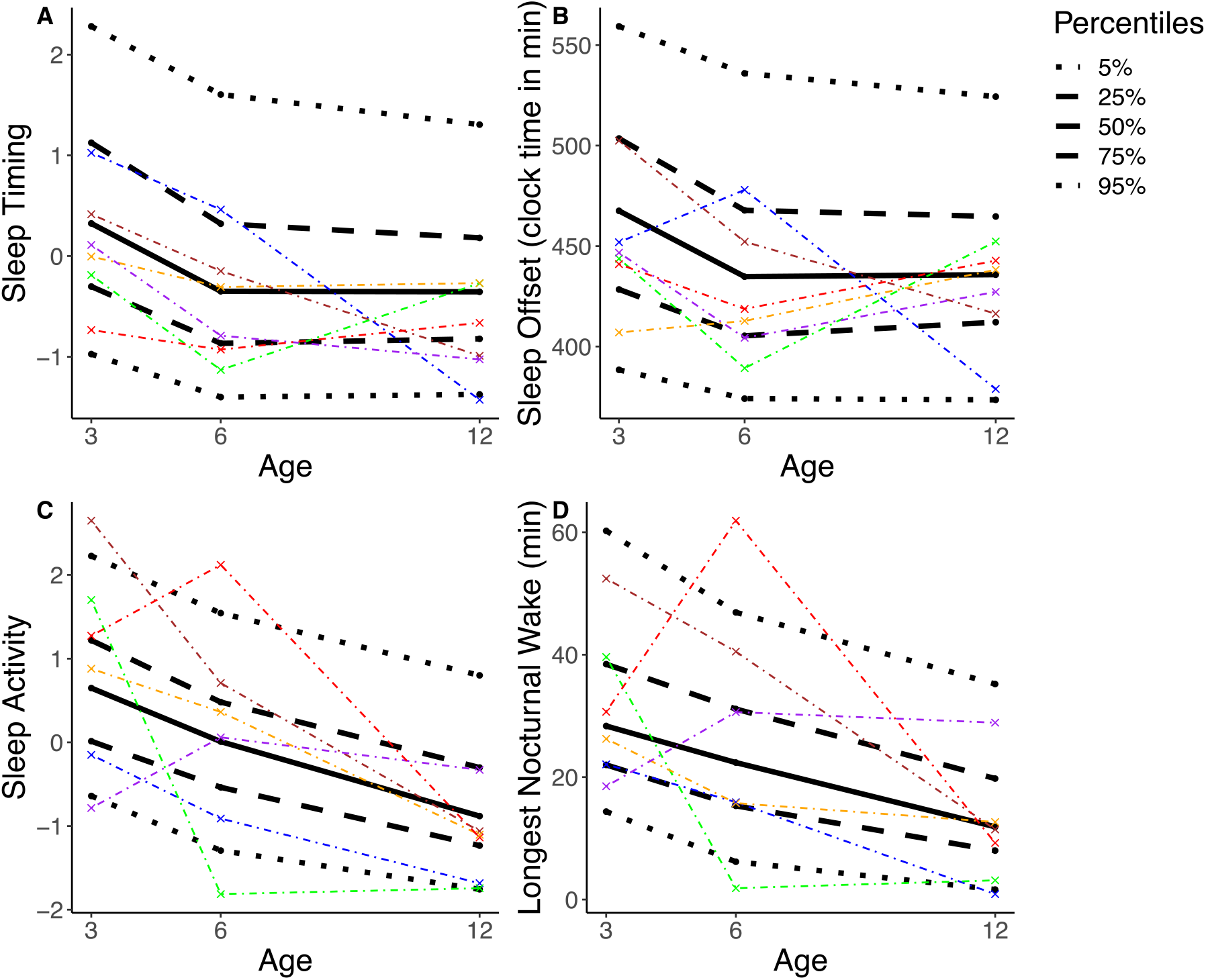
The percentile distribution of two sleep composites *(Sleep Timing, Sleep Activity)* and their highest loading single variable *(Sleep Offset* or *Longest Nocturnal Wake)* is illustrated based on one randomly selected imputation (black; solid line = median, dashed line = interquartile range, dotted line = 90^th^ percentile). 6 randomly selected participants are represented each with specific color. Results illustrate that the position of a participant within the percentile distribution fluctuates across the assessments of 3, 6 and 12 mo. In other words, e.g. an infant with a comparatively high score on *Sleep Activity* at 3 mo does not necessarily maintain a high score in *Sleep Activity* at 6 and 12 mo.

### 3.5 Association of Sleep Composite with Behavioral Development

Lastly, we evaluated whether infant sleep composites are linked to behavioral developmental status. Multilevel models across all assessment timepoints revealed a negative link between *Sleep Day* and ASQ-*Collective score* (b = −6.65, t_(344.65)_ = −2.22, p = 0.03). No association was observed between behavioral development and the other sleep composites (p > 0.05, Table 4). The effect between *Sleep Day* and *Collective Score* was more pronounced after reducing the model to only include *Sleep Day* and control variables (Exact age, sex) and no other sleep composites (b = −7.88, t_(358.69)_ = −2.83, p = 0.005). This association suggests that infants with more daytime sleep had lower overall developmental scores. To investigate this finding in more depth we determined whether the effects persisted in the two behavioral sub-scores *Personal-social* and *Gross Motor.* This was not the case – no significant effects between the behavioral sub-scores and any of the sleep composites were found (p > 0.05). It is thus likely that the effect of *Sleep Day* with behavioral developmental is driven by the combination over multiple scales of development.

**Table 4.** Associations between sleep composites and behavioral development as quantified by the Ages and stages questionnaire. Bold font indicates significant associations (p < 0.05). SE = Standard error of measurement.

| Variable | *Collective Score* |  | *Personal-social* |  | *Gross Motor* |
| --- | --- | --- | --- | --- | --- |
|  | b ± SE | p | b ± SE | p | b ± SE |
| Intercept | 203.16 ± 6.58 | < 0.001 | 42.17 ± 2.04 | < 0.001 | 38.82 ± 2.21 |
| *Sleep Activity* | -0.91 ± 2.27 | 0.69 | -0.46 ± 0.73 | 0.53 | 1.05 ± 0.80 |
| *Sleep Day* | **-6.65 ± 3.00** | **0.03** | -1.08 ± 0.98 | 0.27 | 1.12 ± 1.10 |
| *Sleep Night* | 0.84 ± 2.08 | 0.68 | 0.49 ± 0.62 | 0.43 | 0.68 ± 0.69 |
| *Sleep Timing* | -0.20 ± 2.41 | 0.94 | -0.40 ± 0.70 | 0.57 | 0.47 ± 0.78 |
| *Sleep Variability* | -2.52 ± 2.09 | 0.23 | 0.33 ± 0.67 | 0.62 | 0.55 ± 0.77 |
| Exact age | 0.21 ± 0.79 | 0.78 | -0.37 ± 0.27 | 0.17 | 0.14 ± 0.30 |
| Female sex | 9.49 ± 5.63 | 0.09 | 1.76 ± 1.35 | 0.19 | 0.90 ± 1.55 |

## 4. Discussion

In this study we demonstrate that numerous dimensions of infant sleep can be reduced to the five core sleep composites *Sleep Activity, Sleep Timing, Sleep Day, Sleep Night* and *Sleep Variability*. The sleep composites undergo developmental changes that align with the known maturation of sleep behaviors. We thus recommend the use of sleep composites to reduce variables and to streamline analyses between different lines of research.

Furthermore, both the majority of sleep composites, as well as the single sleep variables, show only limited within-subject stability across the first year of infancy, which contrasts with reports on older children. The only notable exception is *Sleep Timing*, which is stable across the first year of life, and indicates either a parental or infant preference. The lack of within-subject stability can be problematic for studies with a single assessment time-point, because results will vary depending on the assessment time-point. We thus recommend to use multiple assessment time points, especially when the early sleep behavior is used to predict later cognitive or behavioral outcomes. Interestingly, *Sleep Day* is associated with behavioral developmental scores, therefore being a potential marker for maturation. Additionally, we report a sex difference in *Sleep Activity,* with male participants showing more and longer awakenings during the night compared to female infants.

We confirm for the first time the existence of the five infant sleep composites *Sleep Activity, Sleep Timing, Sleep Day, Sleep Night* and *Sleep Variability,* as previously identified in 2.5-3.5 year-old children [27] and which correspond to the most fundamental dimensions of sleep regulation. We adhered to the same terminology used by Staples et al., except for replacing *Sleep Duration* with *Sleep Night* to differentiate it from *Sleep Day.* To represent sleep in the earliest period of life, we included several variables pertaining to daytime sleep (e.g. number of naps, longest duration of consolidated wake). This confirmed *Sleep Day* as a construct separate from the other sleep composites. 14 single sleep variables were excluded from the sleep composites due to low loadings on any of the composites. This included several of the variability variables, as well as *Sleep Latency* which Staples *et al.* also reported to separate from the other composites. In comparison to Staples et al. our method explains sleep variable variance to a slightly lower extent (71% vs 82%), which might be caused by the difference in the assessed sleep variables or the different age range (33 vs 18). Importantly, the proposed sleep composites follow the primary developmental trajectories of the single sleep variables [17,61]. Overall, we conclude that composite sleep dimensions computed from underlying sleep variables are consistent from infancy to childhood and correspond well to the known core maturation of infant sleep patterns.

The selection of variables can be difficult because of the diversity of sleep variables and computations. Using too many variables can lead to multiple testing problems and increase false positive findings [26]. Thus, using composites to reduce the number of variables facilitates investigations in multidimensional research. Our results demonstrate that the resulting sleep composites remain consistent across early development, which aligns with Staples et al., even though different sleep variables were used for computations and even though actigraphy devices differed (GENEActive in our study, MicroMini Motionlogger used by Staples et al.). Our analysis confirms that all single sleep variables identical to Staples et al., [27] loaded onto the same sleep composite. This strongly supports the use of sleep composites, which has the additional advantage to enhance comparability across studies. Moreover, when computation of sleep composites is not possible, our results can guide the selection of variables. Specifically, single sleep variables with high loading are most comparable to the corresponding sleep composites and therefore preferable.

While sleep composites showed some stability across adjacent time periods (3-6 and 6-12 months), the majority of sleep composites did not maintain strong stability across the longest period from 3 to 12 months (except for *Sleep Timing*). This aligns with a previous report, which examined stability of sleep behaviors from 3 to 42 months [11] and found more stability in sleep duration across shorter time intervals while sleep onset time was very stable. Compared to children, adolescents and adults, the stability of sleep variables is exceptionally low in infants. In children 3-7 years old the year-to-year stability was moderate (r = 0.4 – 0.6) in variables related to *Sleep Night* and *Sleep Timing* (even though low stability was noted in *Sleep Activity)* [62]. Thus, the stability of *Sleep Night* increases until childhood, while *Sleep Timing* remains stable and *Sleep Activity* remains variable until adolescence. A 10-year-long study examining dynamics of sleep duration based on interviews from ages 1 to 10 years reported annual fluctuations, yet overall long-term stability [63]. In adults, year-to-year correlation is high for most sleep measures, especially when derived from several nights (r = 0.48 – 0.93) **[64–67]**. While in older children the instability of sleep behaviors might be due to measurement imprecisions and therefore is improved by using composites (shown by Staples et al.), it seems that the instability of sleep behaviors in infancy is inherent in the behavior itself. This is in agreement with the observation that parent-reported infant sleep problems are usually not persistent across longer time periods [68]. Hence, because variability in infant sleep persists naturally, infant sleep composites are not eliminating this variability. One solution to address this point, specifically when examining later outcomes, is to perform a repeated-measures design, as has been previously suggested by Ednick et al. [69]. If this is not possible, it is important to clarify the ages a finding relates to.

The high within-infant stability of *Sleep Timing* is notable and we assume that it is largely parent-driven. This is confirmed by the finding that parent’s bedtimes are positively correlated with *Sleep Timing* (Mother r_s_ = 0.33, p < 0.001 Father r_s_ = 0.24, p < 0.001; exploratory analysis using the reported bedtimes in the Pittsburgh Sleep Quality index). Not surprisingly, parents with later bedtimes also have infants with later sleep timing. However, interestingly, parental bedtimes are not a significant factor in a model that includes within-subject stability of infants to predict infant *Sleep Timing* (Mother F_(1,75156.98)_ = 2.96, p = 0.09, partial η^2^ = 0.01, Father F_(1,60885.42)_ = 2.84, p = 0.09, partial η^2^ = 0.01, participant ID F = 3.33, p < 0.001, partial η^2^ = 0.69). Therefore, variance in infant’s sleep remains unexplained by parent’s bedtime preferences. It is unclear, whether this variance relates to other parental factors (*e.g.* cognitions about regular timing of infant sleep), or if the infants themselves already start to demonstrate daytime preference as an early form of infant chronotype.

Intriguingly, we found a difference in *Sleep Activity* between male and female infants. This is both surprising – because most previous studies in infants reported no sex differences [11,70,71] – and unsurprising – because these differences are well known in adults [72–74]. Boys commonly show higher activity levels [75], which could cause more activity during sleep in infant boys. However, one study reported sex differences as young as age 2 weeks in electroencephalographic recordings [76]. Therefore, with methods that are sufficiently sensitive, sex differences in sleep behaviors can be detected already very early in life.

A final study goal was to examine if any of the sleep composites mirrors behavioral maturation. Thus, we tested the association between sleep composites and behavioral developmental status. Indeed, infants with more daytime sleep (*Sleep Day*) had lower ASQ-*Collective scores.* Our results align with Spruyt et al. who report a negative association between daytime sleep at 12 months with emotional regulation and behavioral maturation [77]. Of relevance might be that the variable *Sleep Day* shows the largest developmental changes across the first year of life. For example, hours asleep during the day and amount of naps are reduced by half from 3 to 12 months of age. Furthermore, the neurophysiology of daytime sleep also changes with age: 5-year old children show decreased slow wave activity (a marker of sleep need) during an afternoon nap compared to 2 and 3-year old children [10], which also suggests that daytime sleep specifically reflects maturation of the central nervous system. When infants are young, napping is important for new memory formation [5,78,79]. When infants get older, their tolerance of longer wake periods increases, which likely also includes their capacity of information acquisition without an immediate nap. Therefore, we hypothesize that a faster decrease in daytime sleep reflects more advanced maturation on a neuronal level. Kurdziel et al. support this theory by demonstrating that naps enhance memory performance in pre-school children only in those children who habitually nap [80] (but see [81]). Therefore, children likely stop to take regular naps when they have developed a physiological tolerance to longer wake periods and when they can retain information without a subsequent nap.

We included a comprehensive list of commonly used sleep variables of infants and young children. Because the structuring of factors in the principle component analysis depends on the variables used, the sleep composites identified in this study might not be representative for other investigations that include different single sleep variables. Furthermore, the choice of 5 factors for the PCA was based on both, data driven criteria, as well as on the interpretability of the resulting sleep composites. It is therefore possible that from a data-driven perspective, more factors would result in a better model fit. However, we prioritized the interpretability of composites so that infant sleep composites can be used for analysis with other datasets. Further, our data is biased towards a higher parental education level (data not shown) and is therefore more homogenous than the general population.

## 5. Conclusions

Our five sleep composites accurately characterize the complex dimensions of infant sleep and reflect known maturational dynamics of infant sleep. To increase comparison across studies, we suggest that researchers use infant sleep composites or, if not possible, single sleep variables with high loadings on the sleep composite of interest. As infant sleep behavior is highly variable both between and within infants, we recommend to use multiple assessment time points, especially for testing sleep behaviors as predictors for later cognitive, emotional or behavioral outcomes. Two exciting future directions of research will likely target *Sleep Timing* as a possible early chronotype and *Sleep Day* as a maturational marker. Therefore, this study opens up new possibilities to standardize and advance the emerging field of infant sleep research.

## Author Contributions

Conceptualization, S.F.S., S.K., R.H. and M.K.; methodology, S.F.S., S.K., R.H. and M.K..; software, S.F.S..; formal analysis, S.F.S..; investigation, S.F.S., S.K..; resources, S.K., R.H. and M.K.; data curation, S.F.S..; writing—original draft preparation, S.F.S.; writing—review and editing, S.F.S., S.K., R.H. and M.K.; visualization, S.F.S..; supervision, S.K.; project administration, S.F.S..; funding acquisition, S.K., S.F.S, R.H.

## Acknowledgments

We thank the parents and infants for participating in our study. We would like to thank the students and interns of the Baby Sleep laboratory for their help with data collection: Sina Boschert-Hennrich, Rita Grolimund, Viktoria Gastens, Lara Barblan, Rahel Nicolet, Melanie Auer, Monika Stoller and Juliane Berger. Additionally, we thank Andjela Markovic, Christophe Mühlematter, Matthieu Beaugrand and Valeria Jaramillo for reviewing code and general discussion and feedback.

## Appendix A

The imputation data set included the participant number (complete), the timepoint of assessment (complete), gender (complete), exact age at assessment (complete or set to assessment timepoint for data points to impute), *Collective Score* (4.5% missing), *Gross Motor* (4.5 % missing) and *Personal Social* (4.5% missing) scores from the Ages and Stages questionnaire [82], gestation age at birth (complete), sleep environment for the baby at night (own room, parent’s room, room shared with sibling; complete for missing data we inferred from the other assessment time points), number of children in family (complete), analysis run for gut microbiota (complete, if there was not gut microbiota data, the run at which the data would have been analyzed was used), probiotics use (yes/no) at 3 (5.2 % missing) and 6 months (3% missing), antibiotics use at 6 (3% missing) and 12 months (6.7% missing, “never”, “unknown time”, “2-4 weeks before assessment”, “< 2 weeks before assessment”), 3 alpha diversity measures for the gut microbiota (Shannon, Observed, Chao all 4.1% missing), gut microbiota cluster (4.1% missing), gut microbiota age prediction (4.1% missing), education mother (0.2% missing) and father (2.8% missing, “none”, “apprenticeship”, “high school”, “university”, “PhD”), bottle and breastfeeding frequency (4.9% missing, 0 for never or rarely breastfed, 1 if breastfed occasionally, regularly or daily). All 48 sleep variables where included, which ranged from 4.9% missing *(Bedtime)* to 22.3% missing *(Variability of Total Sleep Time, Variability of Sleep Efficiency, Variability of Sleep after Wake Onset).* The missing data from each variable was predicted by all other variables in the dataset that correlated with the variable with r ≥ 0.1. The parental education and bottle and breastfeeding frequency variables had to be excluded as predictors because inclusion of them lead to inability to specify the model (see prediction matrix in the supplementary table A2).

